# Fibronectin, DHPS and SLC3A2 Signaling Cooperate to Control Tumor Spheroid Growth, Subcellular eIF5A1/2 Distribution and CDK4/6 Inhibitor Resistance

**DOI:** 10.1101/2023.04.13.536765

**Authors:** Cameron Geller, Joanna Maddela, Ranel Tuplano, Farhana Runa, Yvess Adamian, Robert Güth, Gabriela Ortiz Soto, Luke Tomaneng, Joseph Cantor, Jonathan A. Kelber

## Abstract

Extracellular matrix (ECM) protein expression/deposition within and stiffening of the breast cancer microenvironment facilitates disease progression and correlates with poor patient survival. However, the mechanisms by which ECM components control tumorigenic behaviors and responses to therapeutic intervention remain poorly understood. Fibronectin (FN) is a major ECM protein controlling multiple processes. In this regard, we previously reported that DHPS-dependent hypusination of eIF5A1/2 is necessary for fibronectin-mediated breast cancer metastasis and epithelial to mesenchymal transition (EMT). Here, we explored the clinical significance of an interactome generated using hypusination pathway components and markers of intratumoral heterogeneity. Solute carrier 3A2 (SLC3A2 or CD98hc) stood out as an indicator of poor overall survival among patients with basal-like breast cancers that express elevated levels of DHPS. We subsequently discovered that blockade of DHPS or SLC3A2 reduced triple negative breast cancer (TNBC) spheroid growth. Interestingly, spheroids stimulated with exogenous fibronectin were less sensitive to inhibition of either DHPS or SLC3A2 – an effect that could be abrogated by dual DHPS/SLC3A2 blockade. We further discovered that a subset of TNBC cells responded to fibronectin by increasing cytoplasmic localization of eIF5A1/2. Notably, these fibronectin-induced subcellular localization phenotypes correlated with a G0/G1 cell cycle arrest. Fibronectin-treated TNBC cells responded to dual DHPS/SLC3A2 blockade by shifting eIF5A1/2 localization back to a nucleus-dominant state, suppressing proliferation and further arresting cells in the G2/M phase of the cell cycle. Finally, we observed that dual DHPS/SLC3A2 inhibition increased the sensitivity of both Rb-negative and -positive TNBC cells to the CDK4/6 inhibitor palbociclib. Taken together, these data identify a previously unrecognized mechanism through which extracellular fibronectin controls cancer cell tumorigenicity by modulating subcellular eIF5A1/2 localization and provides prognostic/therapeutic utility for targeting the cooperative DHPS/SLC3A2 signaling axis to improve breast cancer treatment responses.

## INTRODUCTION

Triple-negative breast cancer (TNBC) is marked by the absence of estrogen/progesterone and human epidermal growth factor receptor 2 (HER2 or ErbB2) expression. While there has been some progress made in diagnosing and treating this subtype of breast cancer at early stages, the median survival for TNBC patients who develop or are diagnosed with metastatic disease is a dire 8-10 months, representing a sharp drop from the five-year survival percentages for metastatic breast cancer of other subtypes (∼40%) or non-metastatic TNBC (∼75%) [1; 2]. Thus, there is a critical and urgent need to identify targetable mechanisms that support progression and resistance to therapeutic intervention.

Enrichment of extracellular matrix (ECM) proteins increase the rigidity of solid tumor microenvironments and have been identified as factors contributing to poor patient outcomes and metastasis [3; 4; 5]. Additionally, the relationship between cell-ECM adhesion and cell cycle progression has been previously established [6; 7; 8]. Since inhibitors of cell cycle progression represent an important class of anti-cancer agents with unique utility among TNBC patients [9; 10], understanding how ECM proteins affect TNBC cell plasticity and tumorigenicity is critical to advance efforts aimed at improving patient outcomes. In this regard, previous work from our group demonstrated that activating the eukaryotic initiation factors 5A 1/2 (eIF5A1 and eIF5A2) via post-translational hypusination drives the production of the PEAK1 (pseudopodium-enriched atypical kinase 1) pseudokinase [11] and that GC7 blockade of DHPS-dependent hypusination abrogated fibronectin-/TGFβ-induced lung metastasis of TNBC cells [12].

Importantly, eIF5A1/2 are currently the only known targets of the two-step hypusination reaction involving spermidine and catalyzed by deoxyhypusine synthase (DHPS) and deoxyhypusine hydroxylase (DOHH) [13]. Previous work has identified a role for eIF5A1/2 during cancer progression [14] and proteomic profiling of TNBC patient samples revealed a positive association between eIF5A2 expression and inhibitors of apoptosis [15]. Moreover, eIF5A1/2 hypusination is detectable across a panel of TNBC cells and can be significantly reduced by the DHPS inhibitor GC7 (N1-guanyl-1,7-diamineoheptane) [12; 16]. Importantly, GC7 inhibits cell viability across these same TNBC cell models [12], and Liu and colleagues previously reported that TNBC cells were sensitized to doxyrubicin-induced cell death by GC7 or siRNA inhibition of DHPS or eIF5A2, respectively [17]. In addition to these cytoplasmic translation functions of eIF5A1/2, recent work has uncovered roles for hypusination-dependent nucleocytoplasmic transport of eIF5A1/2 in the export of post-transcriptionally processed mRNAs [18].

Here, we sought to identify eIF5A1/2- and heterogeneity-related markers that might cooperate and provide novel targetable mechanisms of TNBC progression. Since we discovered that solute carrier 3A2 (SLC3A2 or CD98hc) predicts poor patient survival in basal breast cancers with elevated levels of DHPS, and both DHPS and SLC3A2 have previously established roles in determining the biological outcomes of fibronectin signaling [12; 19] and can regulate proliferation [20; 21], we sought to evaluate how dual inhibition of these proteins affects fibronectin function and TNBC tumorigenicity. The data presented here establish previously unreported roles for fibronectin in TNBC spheroid growth, subcellular transport of the translation factors eIF5A1/2, cell cycle arrest and enrichment of tumorigenic cell subpopulations. Notably, we discovered that these effects of fibronectin on TNBC cells can be blocked by dual DHPS/SLC3A2 inhibition and that this targeting approach represents a unique approach for rendering Rb-negative TNBC cells sensitive to CDK4/6 inhibitors. Thus, we identify a new ECM-dependent mechanism by which the tumorigenicity of TNBC cells is exacerbated and provide data to support this mechanism as a targetable means for improving the efficacy anti-cancer therapies.

## RESULTS

### Identification of DHPS and SLC3A2 upregulation as prognostic factors in basal-like breast cancers

Research over the past decade has uncovered a functional connection between increased cell state heterogeneity and improved fitness of malignant cells during progression of solid tumor types [22]. To better understand the role of eIF5A1/2 in tumor cell plasticity and intratumoral heterogeneity, we developed a bioinformatics pipeline (Figure 1) that used a set of literature-curated [13; 23; 24; 25; 26; 27; 28], cancer-associated heterogeneity (51 genes) and eIF5A1/2 pathway (12 genes) markers (Figure 1A) as input data. Using this 63 gene list, we used the Cytoscape Agilent Literature Search tool to generate a 32-node interactome representing known biochemical and functional eIF5A1 and eIF5A2 connections (Figure 1B). Using the DAVID database to search for ontologies represented among these 32 genes, we identified notable enrichments associated with cell cycle regulation and cell proliferation (Figure 1C).

**Figure 1:**
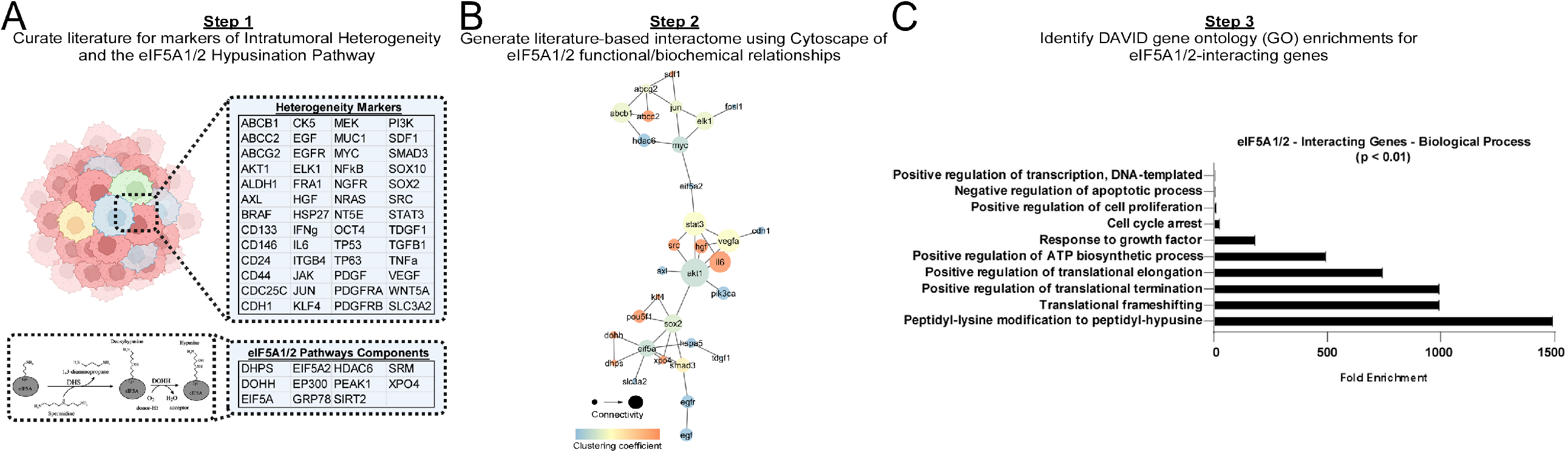
Bioinformatic analyses of intratumoral heterogeneity and eIF5A1/2 pathway markers. A. Literature-mined heterogeneity and eIF5A1/2 pathway markers used for bioinformatics analysis. B. Cytoscape-generated interactome of eIF5A1/2 pathway markers and associated intratumoral heterogeneity markers. “Connectivity” refers to the number of connections a single gene shares with other genes in the interactome; higher connection numbers yields larger node symbol. “Clustering Coefficient” refers to the degree to which an interactome node clusters together with other nodes. Node colors indicate clustering coefficient values. C. DAVID database gene ontology enrichment for biological processes within the set of 11 eIF5A1/2-Interacting genes from (B).

While nearly all direct eIF5A1/2-interacting genes from the heterogeneity interactome were expressed in both normal breast and breast cancer tissues (Figure S1), we noted that eIF5A-dependent translation has been previously reported to support solute carrier 3A2 (SLC3A2, CD98 heavy chain) protein expression in cancer cells [29] and SLC3A2 represents a unique cell surface target that has previously established roles in cell proliferation and tumor progression [21; 30]. Additionally, SLC3A2 mediates polyamine uptake – an established prerequisite for eIF5A1/2 hyusination/activation and tumor progression [13; 31]. Given the previously established reciprocal relationship between eIF5A1/2 and solute carrier 3A2 (SLC3A2 or CD98 heavy chain) signaling [29; 32], that SLC3A2 regulates fibronectin-integrin signaling [19; 33; 34] and that SLC3A2 is a prognostic factor in highly proliferative breast cancer subtypes [35], we evaluated the relationships between expression levels of DHPS, DOHH, eIF5A or eIF5A2 in combination with SLC3A2 across the four primary subtypes of breast cancer (i.e., luminal A, luminal B, HER2-positive and basal). In agreement with previous work showing that SLC3A2 may serve as a strong target candidate in triple-negative breast cancers [36], we discovered that basal breast cancer patients with elevated expression levels of both DHPS and SLC3A2 exhibited significantly decreased overall survival (Figure 2A).

**Figure 2:**
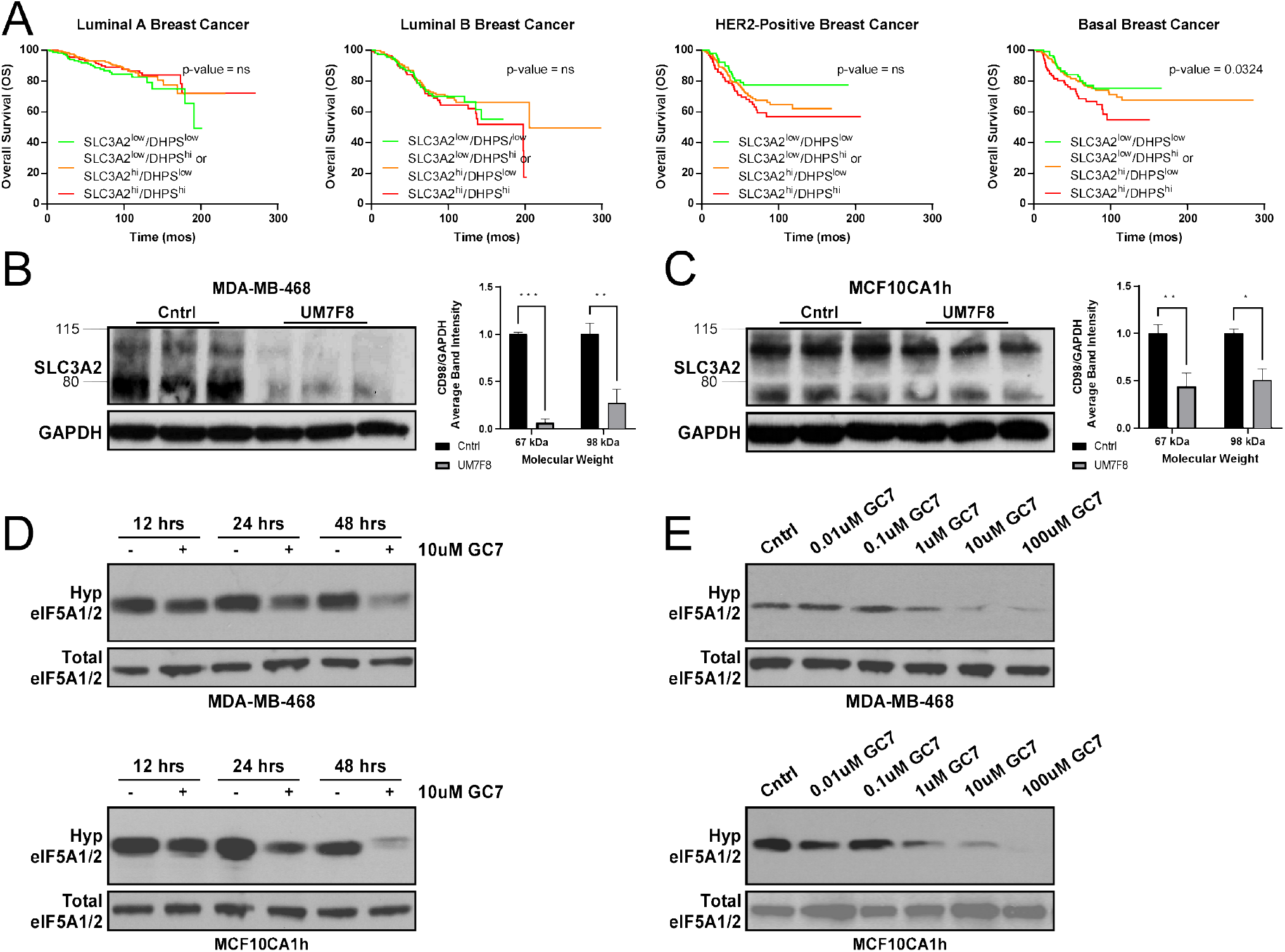
Identification of DHPS and SLC3A2 upregulation as a prognostic factor in basal-like breast cancers. A. Kaplan-Meier plots and Log-Rank analyses for overall survival of breast cancer patients across PAM50 classifications and subdivided by combinations of DHPS and SLC3A2 expression levels. B-C. Western blot analysis for SLC3A2 expression in response to control and 250 ng/mL SLC3A2-targeting monoclonal antibody (UM7F8) in MDA-MB-468 (B) and MCF10CA1h (C) cells. D. Western blot analysis for hypusinated and total eIF5A1/2 levels in response to control or 10uM GC7 treatment of MDA-MB-468 (top) and MCF10CA1h (bottom) cells at indicated time points. E. Western blot analysis for hypusinated and total eIF5A1/2 levels in response to 48 hour control or dose-response GC7 treatment of MDA-MB-468 (top) and MCF10CA1h (bottom) cells. *, **, *** or **** indicate p-values of 0.05, 0.01, 0.001 or 0.001, respectively.

Thus, we tested a previously reported antibody-induced ubiquitin ligase-mediated SLC3A2 downregulation protocol in triple negative breast cancer (TNBC) cells that employs a mouse monoclonal anti-human SLC3A2 antibody (clone UM7F8, IgG1) combined with a goat anti-mouse IgG F(ab′)2 antibody [37]. This approach achieved a significant UM7F8-dependent SLC3A2 downregulation in MDA-MB-468 and MCF10CA1h cell models of TNBC (Figure 2B-C). Concurrently, we verified that the spermidine analog GC7 (N1-guanyl-1,7-diamineoheptane) was able to inhibit eIF5A1/2 hypusination in a time- and dose-dependent manner across both the MDA-MB-468 and MCF10CA1h cells lines (Figures 2D and E).

### DHPS and SLC3A2 cooperate with fibronectin to govern breast cancer spheroid growth

We leveraged these DHPS and SLC3A2 inhibition strategies to evaluate how these pathways affect three-dimensional (3D) spheroid growth using MDA-MB-468 and MCF10CA1h cells. As shown in Figure 3A, MDA-MB-468 cells form well-established single 3D spheroids in ultra-low attachment (ULA) round bottom 96-well plates over the course of two days that can be imaged and quantified using the live-cell IncuCyte platform. We observed that inhibition of DHPS or SLC3A2 at effective doses identified in Figure 2 significantly reduced MDA-MB-468 spheroid growth (Figures 3A-C). Notably, blocking both DHPS and SLC3A2 together decreased spheroid growth more substantially than inhibition of either target by itself. Since SLC3A2 has dual functions as a core constituent of the CD98 transporter and as a regulator of extracellular matrix (ECM) protein-integrin mechanosignaling [38; 39; 40], and we have previously reported that DHPS-mediates TGFβ/fibronectin-induced EMT and metastasis in TNBC [12], we asked whether exogenous fibronectin added into 3D spheroid cultures can alter TNBC cell response to DHPS and/or SLC3A2 inhibition. Interestingly, fibronectin-spiked spheroid culture media could oppose the spheroid-reducing effects of either inhibitor by itself or when delivered together, though dual DHPS/SLC3A2 blockade still limited MDA-MB-468 spheroid growth substantially more than either inhibitor by itself (Figures 3B-C). In contrast, MCF10CA1h cells formed several smaller spheroids under these same conditions that expanded less substantially over the same two-day time period (Figure 3D). Nonetheless, applying a similar spheroid quantification approach revealed that DHPS or SLC3A2 inhibition significantly limited spheroid growth in cultures without fibronectin, and dual inhibition yielded the smallest spheroid growth margins (Figures 3D-F). Interestingly, the addition of exogenous fibronectin into MCF10CA1h spheroid cultures significantly increased spheroid size and rescued the inhibitory effect of DHPS inhibition - an effect could be most efficiently blocked by dual DHPS/SLC3A2 inhibition (Figure 3F).

**Figure 3:**
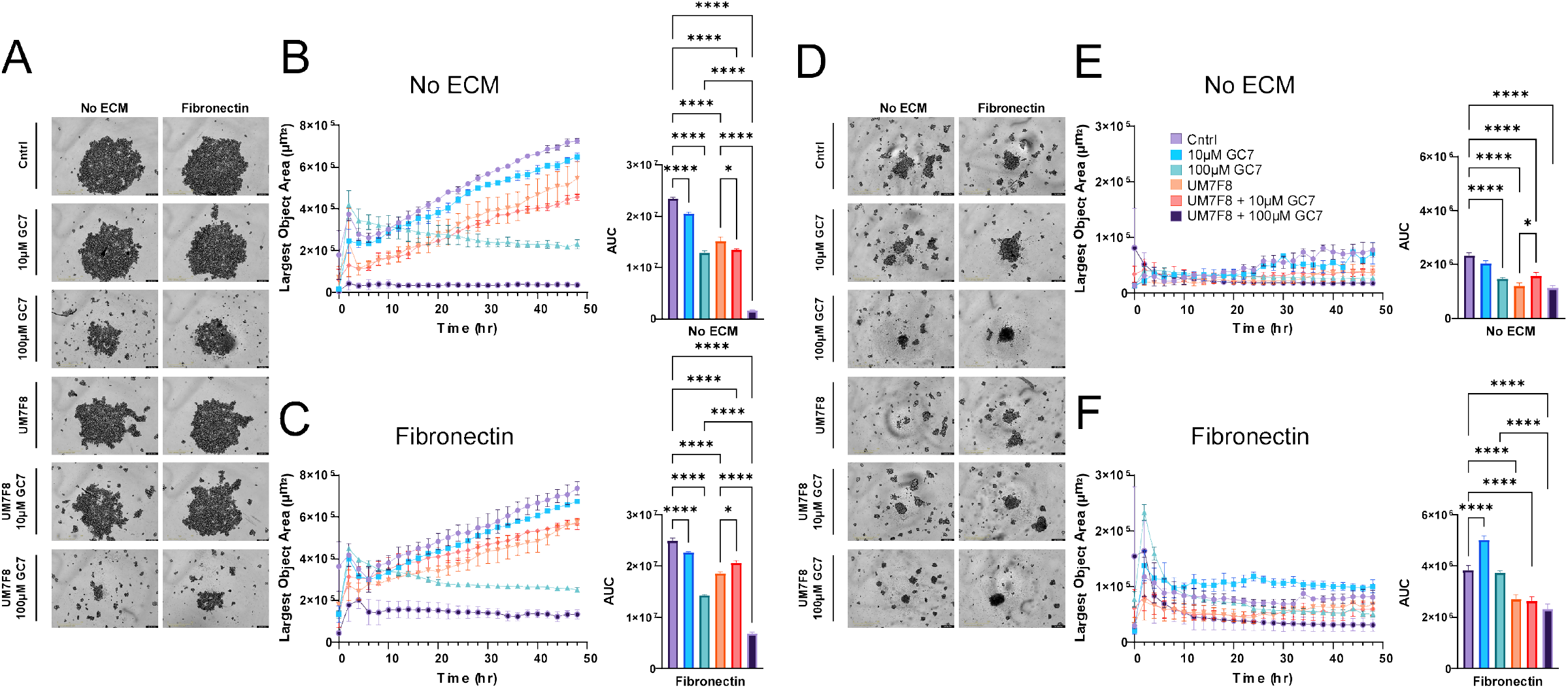
DHPS and SLC3A2 cooperate with fibronectin to govern breast cancer spheroid growth. A. Representative images of MDA-MB-468 cells cultured as spheroids for 48 hours in ULA round-bottom plates using defined media without (left) or with (right) 10 ug/mL fibronectin and in the presence of indicated doses of GC7 and/or 250 ng/mL UM7F8. B-C. Quantification of largest object area for time-lapse imaging of cells in (A) without (B) and with (C) fibronectin. D. Representative images of MCF10CA1h cells cultured as spheroids for 48 hours in ULA round-bottom plates using defined media without (left) or with (right) 10 ug/mL fibronectin and in the presence of indicated doses of GC7 and/or 250 ng/mL UM7F8. E-F. Quantification of largest object area for time-lapse imaging of cells in (D) without (E) and with (F) fibronectin. *, **, *** or **** indicate p-values of 0.05, 0.01, 0.001 or 0.001, respectively.

### Fibronectin induces DHPS-/SLC3A2-dependent cytoplasmic localization of eIF5A1/2 in TNBC cells

While DHPS-dependent hypusination controls the function of eIF5A1/2 in both cytoplasmic [41] and nuclear [42] compartments, the fully hypusinated forms of eIF5A1/2 reside predominantly in the cytoplasm [43; 44]. Previous work by Asku and colleagues identified exportin 4 (XPO4) as a common factor mediating nucleocytoplasmic shuttling of both eIF5A1/2 [18]. Since fibronectin/TGFβ stimulation of TNBC cells enhances eIF5A1/2 hypusination [12] and others have reported a role for ECM-integrin mechanosignaling in the control of subcellular distribution for factors like Yes Associated Protein (YAP) and Rho GTPases [45], we sought to evaluate whether DHPS, SLC3A2 and/or fibronectin control subcellular localization of eIF5A1/2 within TNBC cells.

Interestingly, inhibition of either DHPS or SLC3A2 in either MDA-MB-468 or MCF10CA1h cells cultured in the absence of exogenous fibronectin shifted eIF5A1/2 localization from predominantly nuclear toward a more even nucleocytoplasmic distribution (Figures 4A and B, top). When the DHPS and SLC3A2 inhibitor treatments were combined under these same culture conditions, eIF5A1/2 showed the strongest shift toward this balanced nucleocytoplasmic distribution (Figures 4A and B, top). Generally, the distribution of cells in terms of their subcellular eIF5A1/2 localization patterns was Gaussian. Notably, we observed that exogenous fibronectin induced a strong cytoplasmic eIF5A1/2 localization effect and bimodal distribution pattern in a subpopulation of MDA-MB-468 cells (Figures 4A, bottom). In contrast, exogenous fibronectin had little effect on eIF5A1/2 localization patterns in MCF10CA1h cells (Figures 4B, bottom). Interestingly, DHPS or SLC3A2 inhibition opposed the effect of exogenous fibronectin and shifted eIF5A1/2 localization back into the nucleus of some cells in both MDA-MB468 (Figure 4A, bottom) and MCF10CA1h (Figure 4B, bottom) cells. However, only dual DHPS/SLC3A2 inhibition was effective at fully reversing the fibronectin-induced cytoplasmic localization of eIF5A1/2 to the cytoplasm (Figures 4A and B, bottom) and recovering the Gaussian and nuclear distribution of eIF5A1/2 localization in MDA-MB-468 cells (Figure 4A, bottom).

**Figure 4:**
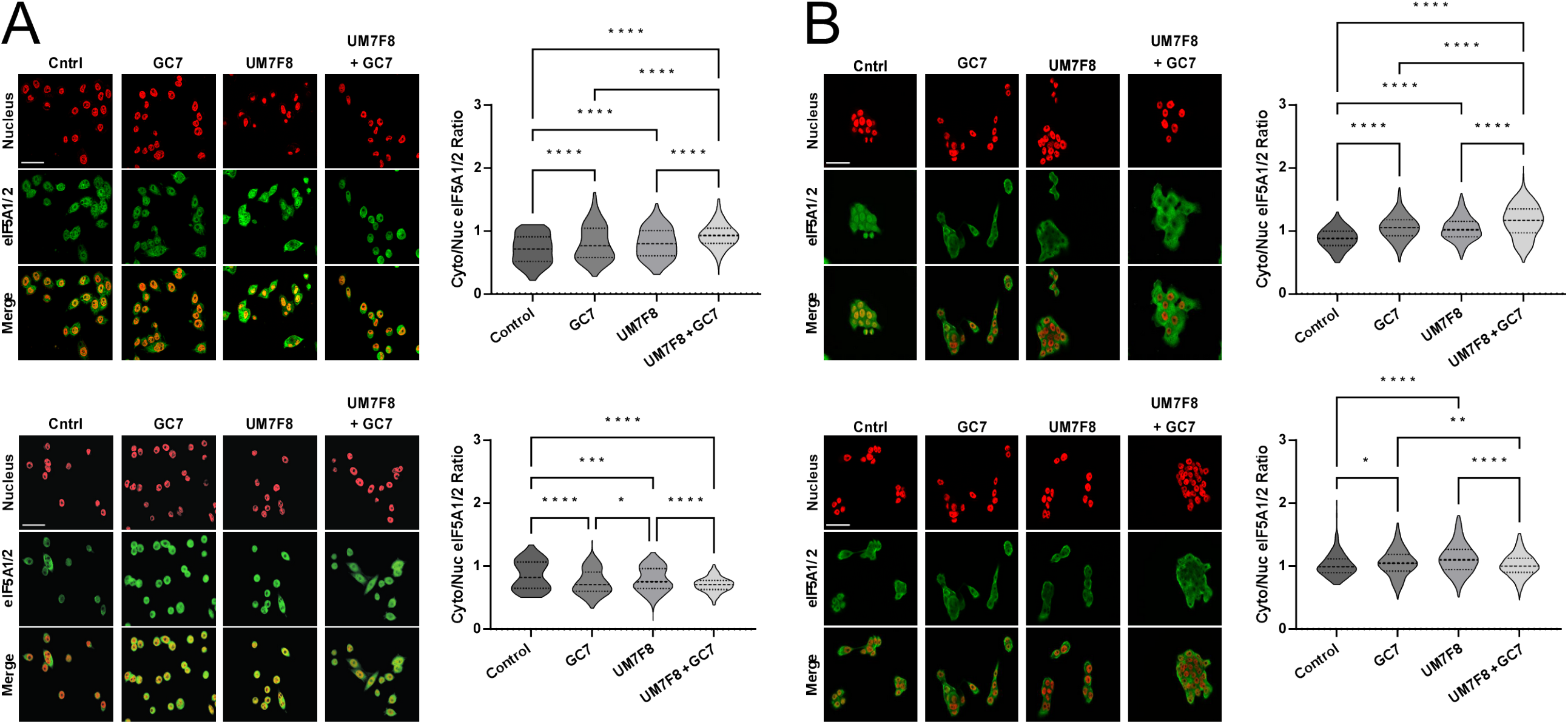
Fibronectin induces DHPS-/SLC3A2-dependent cytoplasmic localization of eIF5A1/2 in TNBC cells. A-B. Representative images (left) and quantification (right) of the cytoplasmic:nuclear ratio for eIF5A1/2 in MDA-MB-468 (A) or MCF10CA1h (B) cells plated on no extracellular matrix (ECM, top) or on 5 ug/mL fibronectin (bottom) and treated with control or 10 uM GC7 and/or 250 ng/mL UM7F8 for 48 hours. Scale bars represent 50 um. *, **, *** or **** indicate p-values of 0.05, 0.01, 0.001 or 0.001, respectively.

### Fibronectin induces G0/G1 arrest and sensitizes TNBC cells to a further G2/M arrest following dual DHPS/SLC3A2 inhibition

Previous work has connected mechanoresponsive integrin signaling with cell cycle control [6; 7; 46]. Thus, we sought to evaluate how fibronectin-based integrin signaling affected the cell cycle profiles of MDA-MB-468 and MCF10CA1h cells (Supplemental Figure 2). Interestingly, while fibronectin did not affect the population of MDA-MB-468 cells within the proliferative S-phase, it markedly increased the percentage of cells in G0/G1 phase and decreased cells in the G2/M phase, consistent with these cells continuing to progress through the cycle but with reduced activity of the G0/G1 checkpoint regulators (Figures 5A and B). Like fibronectin, treating MDA-MB-468 cells with inhibitors of SLC3A2 or both DHPS and SLC3A2 caused a cell cycle arrest in the G0/G1 phase in the absence of exogenous fibronectin. Additionally, dual DHPS/SLC3A2 inhibition also caused a significant reduction in MDA-MB-468 cells within the proliferative S-phase (Figures 5A and B). In contrast, treating MDA-MB-468 cells with DHPS and SLC3A2 inhibitors in the presence of exogenous fibronectin caused cells to arrest in the G2/M phase. While inhibition of DHPS and/or SLC3A2 did not affect the cell cycle profile of MCF10CA1h cells in the absence of exogenous fibronectin, DHPS and/or SLC3A2 inhibition caused fibronectin-treated cells to arrest in the G2/M phase (Figures 5C and D).

**Figure 5:**
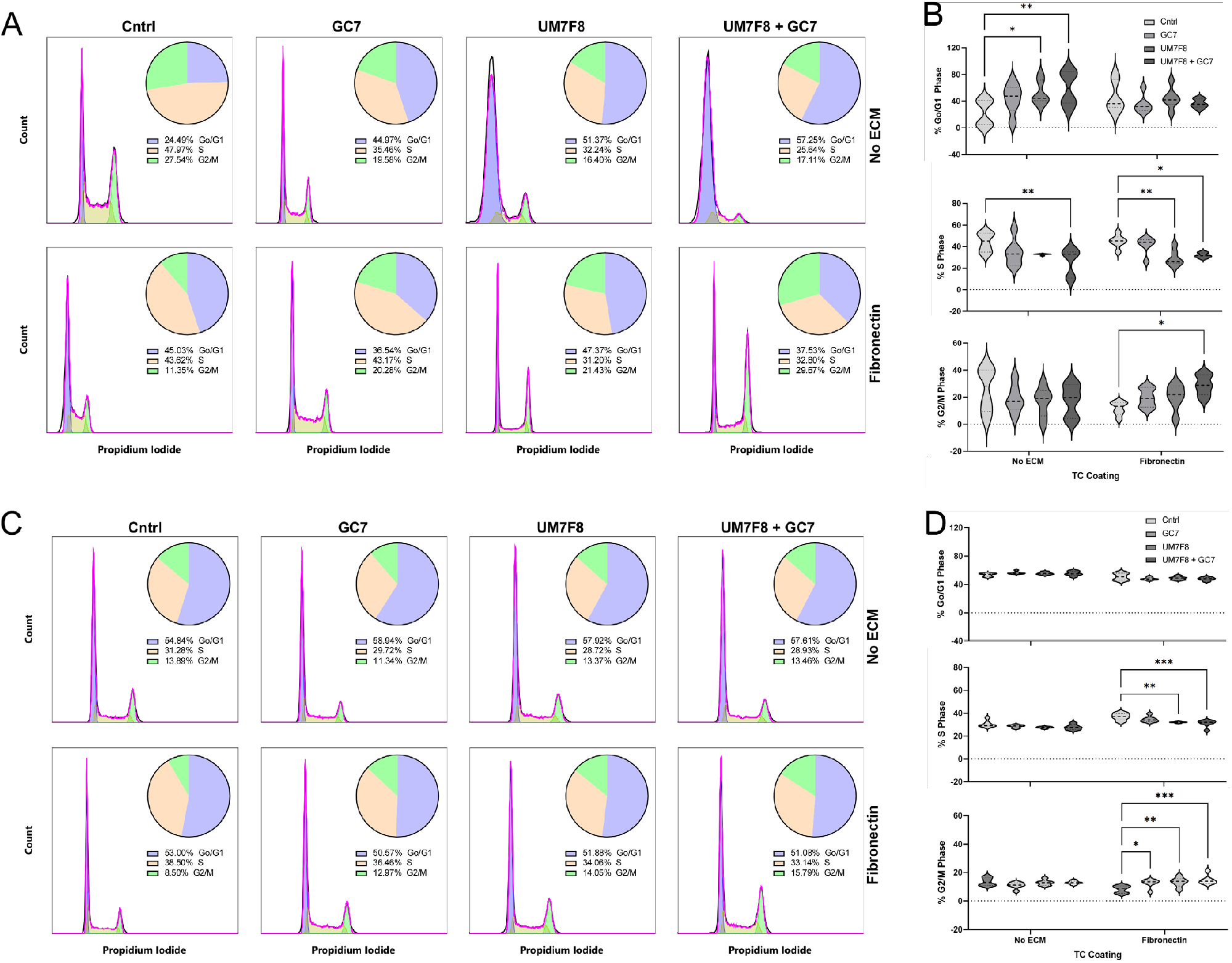
Fibronectin imposes a G0/G1 arrest on TNBC cells and sensitizes them to a G2/M phase arrest by dual DHPS/SLC3A2 inhibition. A-B. Cell cycle plots and associated phase percentages (pie graph inlays) (A) and quantification (B) for MDA-MB-468 cells plated on no extracellular matrix (ECM, top or left) or on 10 ug/mL fibronectin (bottom or right) and treated with control or 10 uM GC7 and/or 250 ng/mL UM7F8 for 48 hours. C-D. Cell cycle plots and associated phase percentages (pie graph inlays) (C) and quantification (D) for MCF10CA1h cells plated on no extracellular matrix (ECM, top or left) or on 10 ug/mL fibronectin (bottom or right) and treated with control or 10 uM GC7 and/or 250 ng/mL UM7F8 for 48 hours. *, **, *** or **** indicate p-values of 0.05, 0.01, 0.001 or 0.001, respectively.

### Dual DHPS/SLC3A2 inhibition sensitizes TNBC cells to CDK4/6 inhibition

Our observations that fibronectin arrests TNBC cells in the G0/G1 cell cycle phase (Figure 5) and that dual DHPS/SLC3A2 in TNBC cells exposed to fibronectin further arrested cells in the G2/M phase (Figure 5) led us to evaluate the effects of DHPS and/or SLC3A2 inhibition on TNBC cell response to CDK4/6 inhibition. Notably, MDA-MB-468 cells (in contrast to MCF10CA1h cells) lack expression of the Rb protein – a critical determinant in tumor cell responsiveness to CDK4/6 inhibitors. In agreement with this, treatment of these cells with Palbociclib had no effect on cell growth curves (Figures 6A and B). Interestingly, MDA-MB-468 cells grown on fibronectin substrates grew faster when treated with Palbociclib (Figures 6A and C). In contrast to 3D spheroid cultures (Figure 3) and in agreement with our previous work [12], MDA-MB-468 cells were not sensitive to DHPS inhibition alone when grown as 2D cultures in the absence of fibronectin (Figures 6A – C). Similarly, DHPS and/or SLC3A2 inhibition elicited only modest effects on MDA-MB-468 cell growth in the absence or presence of fibronectin (Figures 6A – C). Interestingly, however, MDA-MB-468 cells grown on fibronectin substrates (i.e., cells with increased cytoplasmic eIF5A1/2 levels and arrested in the G0/G1 phase of the cell cycle), became sensitive to pablociclib-induced cytostasis when DHPS and/or SLC3A2 inhibitors were applied (Figures 6A and C). In contrast to the CDK4/6 inhibitor-resistant MDA-MB-468 cells, the growth curves of MCF10CA1h cells grown in the absence or presence of fibronectin were suppressed by Palbociclib treatment (Figures 6D – F). Importantly, however, dual inhibition of DHPS and SLC3A2 further sensitized these cells to Palbociclib-indued cytostasis (Figures 6D – F).

**Figure 6:**
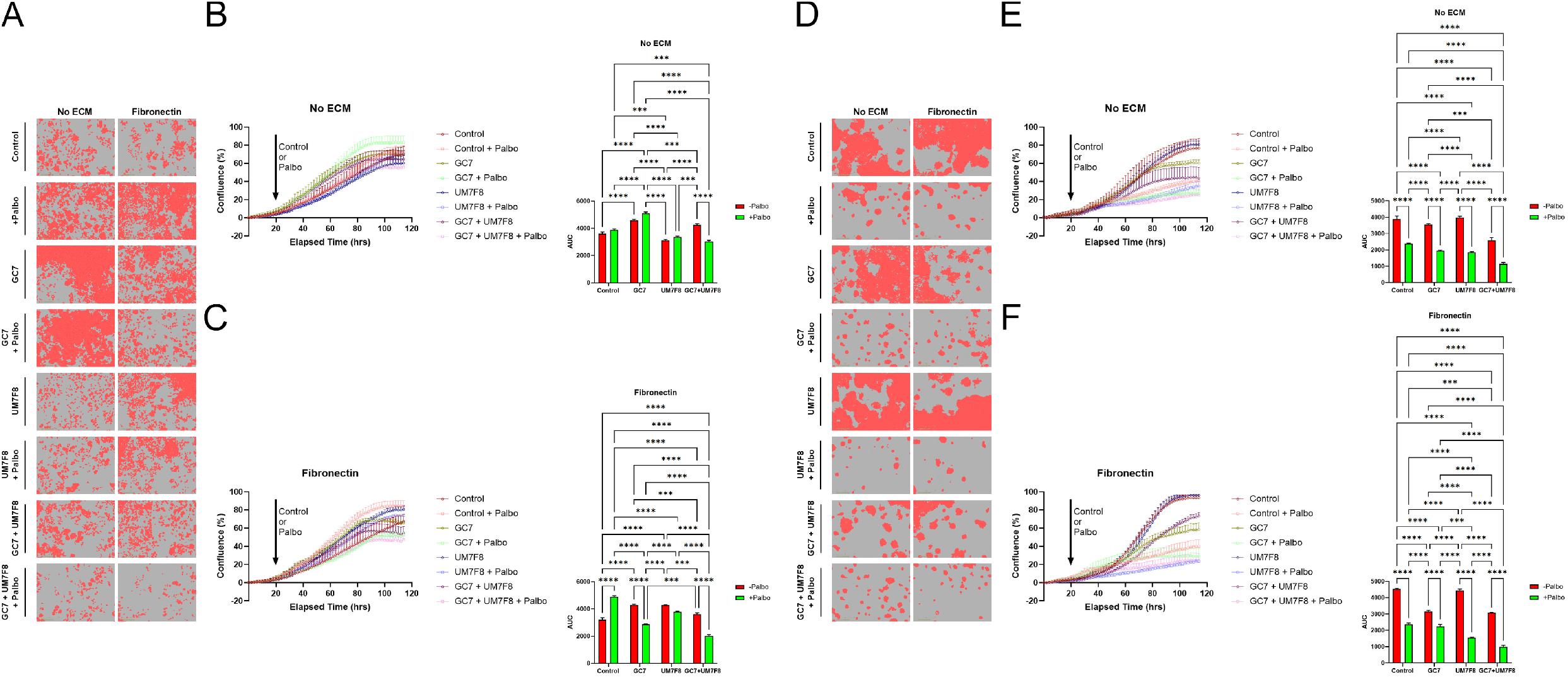
Dual DHPS/SLC3A2 inhibition sensitizes TNBC cells to CDK4/6 inhibition. A. Representative images of MDA-MB-468 cells (and quantification masks) after 80 hours of growth in 96-well plates. Cells were pretreated with DHPS and/or SLC3A2 inhibitors for 48 hours before plating and retreated upon plating into wells without (No ECM) or with 10 ug/mL fibronectin (Fibronectin). Control or Palbociclib treatments were added 24 hours after plating. B-C. Quantification of cell confluence area across time-lapse imaging of cells in (A) without (B) and with (C) fibronectin. D. Representative images of MDA-MB-468 cells (and quantification masks) after 80 hours of growth in 96-well plates. Cells were pretreated with DHPS and/or SLC3A2 inhibitors for 48 hours before plating and retreated upon plating into wells without (No ECM) or with 10 ug/mL fibronectin (Fibronectin). Control or Palbociclib treatments were added 24 hours after plating. E-F. Quantification of cell confluence area for time-lapse imaging of cells in (D) without (E) and with (F) fibronectin. *, **, *** or **** indicate p-values of 0.05, 0.01, 0.001 or 0.001, respectively.

### Cooperative DHPS/SLC3A2 function and signaling in TNBC

The proposed model in Figure 7 depicts the functional interplay of cell surface SLC3A2 with intracellular DHPS activities. Extracellular fibronectin shifts eIF5A1/2 localization from nuclear to cytoplasmic, correlating with an increased sensitivity to dual DHPS and SLC3A2 inhibition. Under these inhibitory conditions, cells are arrested in both the G0/G1 and G2/M phases of the cell cycle and sensitized to the CDK4/6 inhibitor Palbociclib – even in cells lacking the standard CDK4/6 inhibitor biomarker, Rb.

**Figure 7:**
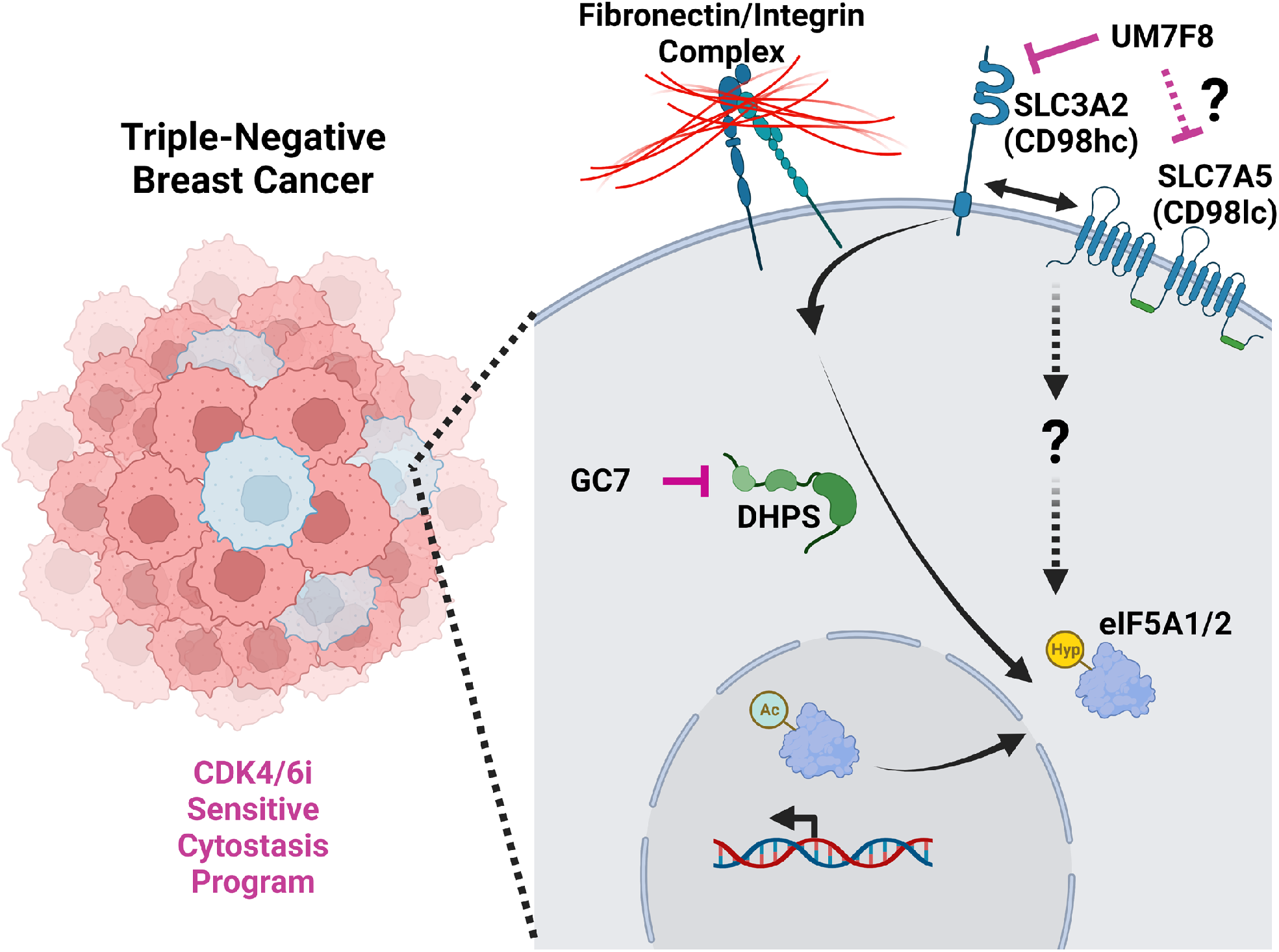
Model of cooperative DHPS/SLC3A2 function in TNBC. SLC3A2 and DHPS are co-expressed in triple-negative breast cancers and predict poor survival. Fibronectin can induce localization of eIF5A1/2 to the cytoplasm. Importantly, these subcellular effects of fibronectin are mediated by DHPS and SLC3A2 function and confer DHPS-/SLC3A2-dependent cytostasis that can be targeted to sensitize cells to CDK4/6 inhibitors.

## DISCUSSION

Here, we report a previously unknown functional relationship between fibronectin, DHPS and SLC3A2 in the context of triple-negative breast cancer (TNBC) tumorigenesis and therapy response. Hypusine and the hypusination pathway, that impinges upon the activity of both eIF5A1 and eIF5A2, are known to be critical regulators of eukaryotic cell translation [28]. Specifically, active/hypusinated eIF5A1/2 have been demonstrated to play several roles in protein translation – i.e., stimulation of peptide bond formation at consecutive proline sequences [47], reduction of ribosome-stalled sites [48] and enhancement of translation termination [49]. Thus, understanding the extracellular and microenvironmental factors that impact this pathway holds promise for leveraging normal cellular physiology and pathophysiological processes to abrogate cancer progression.

In this regard, we previously identified that eIF5A1/2 hypusination is required for translation of PEAK1 (a polyproline-containing protein) [11] and that PEAK1 mediates TGFβ/fibronectin-induced EMT and metastasis of TNBC cells [50]. While TGFβ signaling does not alter total cellular levels of eIF5A1/2, we recently reported that it can increase levels of hypusinated eIF5A1/2 protein in TNBC cells – an effect that correlated with DHPS-dependent metastatic dissemination of TNBC cells [12]. It has been shown that eIF5A1/2 are translocated into the cytoplasm in response to hypusination [43]. However, these previous studies depended on overexpression of eIF5A1/2 mutants with altered lysine 48 and 50 residues (K48A/K50A) as a means to control hypusination. The work presented here is the first to describe this subcellular translocation phenomenon for endogenous eIF5A1/2 and in response to extracellular fibronectin stimulation (Figure 4). Our observation that fibronectin can control translation factor function is in agreement with previous work by Gorrini and colleagues demonstrating that fibronectin controls cap-depednent translation through integrin beta 1 (ITGB1) [51]. Nonetheless, the spatiotemporal dynamics of eIF5A1/2 subcellular trafficking and whether combining TGFβ with fibronectin potentiates cytoplasmic localization of eIF5A1/2 remain to be defined. Furthermore, it will be relevant for future work to assess the cooperative role of DHPS and SLC3A2 in fibronectin/PEAK1-dependent switching of TGFβ signaling outcomes from predominantly Smad2/3-dependent tumor suppressive to non-canonical MAPK-dependent invasion/metastasis promoting.

The other primary beta integrin subunit to mediate fibronectin signaling is integrin beta 3 (ITGB3) [52]. ITGB3 has been previously demonstrated to regulate cellular senescence by activating autocrine and paracrine TGFβ signaling [53]. The data we present here (Figure 5) suggest that fibronectin can also act independently of TGFβ to reduce cell cycle progression through the G0/G1 phase and increase cytoplasmic eIF5A1/2 localization. Though, as discussed above, the cooperativity of exogenous TGFβ and fibronectin in this system remains to be explored. Importantly, we show that only when cells are stimulated with fibronectin does dual DHPS/SLC3A2 inhibition prevent cell proliferation and induce a complete cell cycle arrest suggesting that this fibronectin-induced cytoplasmic eIF5A1/2 localization represents an important targetable vulnerability that may be leveraged to suppress solid tumor progression (Figure 5). In the context of targeted cell cycle inhibitors for treating solid tumor progression, CDK4/6 serine/threonine kinases have emerged as important candidates [54]. As with many promising targeted therapeutics, patients often exibit intrinsic or acquired resistance – factors predicting poor disease outcomes. The primary biomarker for intrinsic resistance to CDK4/6 inhibition is the loss of the Retinoblastoma (Rb) gene. Mechanistically, this is due to the ability of CDK4/6 to phosphorylate Rb and for phosphorylated Rb to inhibit E2F-dependent transcription of proliferation-driving genes such as CDK1/2/3 and their respective cyclins [54]. However, other mechanisms – even in the absence of Rb proteins – can contribute to CDK4/6 inhibitor resistance. For example, Yoshida and colleagues recently reported that Rb loss can drive the expression of SLC36A1 via increased E2F-dependent transcription [55]. While SLC3A2 was not tested in this study, they reported that expression of the SLC3A2 amino acid transporting co-receptor SLC7A5 (CD98 light chain) was decreased in the Rb-negative melanoma cell variants response to CDK4/6 inhibition. Furthermore, in these Rb-negative variants, targeting the mTORC1 pathway presented a viable mechanism to overcome CDK4/6 inhibitor resistance in the absence of Rb. Here, we describe a unique fibronectin/DHPS/SLC3A2 signaling axis that, when targeted, renders Rb-negative MDA-MB-468 TNBC cells sensitive to the CDK4/6 inhibitor Palbociclib (Figure 6).

While SLC3A2 has been previously reported to affect cell proliferation and adhesion across multiple cell types via bimodal amino acid transport and fibronectin/integrin signaling mechanisms [38; 56], the work presented here underscores the importance of deciphering the respective contributions of these different mechanisms of action for SLC3A2 in the context of solid tumor biology. In this context, it is also relevant to consider the contribution of fibronectin/integrin signaling in cell cycle control [57]. For example, it is possible that the fibronectin/integrin and amino acid regulatory functions of SLC3A2 cooperate to stimulate eIF5A1/2 activity and cytoplasmic localization. However, our findings (Figure 4) showing a differential response of TNBC cells to SLC3A2 inhibition, in terms of eIF5A1/2 localization, depending upon whether the cells are exposed to extracellular fibronectin. This suggests that these biomodal functions of SLC3A2 may oppose one another.

Taken together, the findings reported here uncover a new DHPS/SLC3A2 mechanism that controls eIF5A1/2 activity and is stimulated by extracellular fibronectin – laying the foundation for preclinical studies aimed at evaluating the efficacy of dual DHPS/SLC3A2 targeting strategies to sensitize TNBC and other types of solid tumors to CDK4/6 inhibitors and/or other targeted therapies.

## METHODS

### Cell culture

MCF10ACa1h cells were purchased from the Karmanos Cancer Institute (Detroit, MI). MDA-MB-468 cells were purchased from the American Type Culture Collection (ATCC; Manassas, VA). All cells were cultured according to source instructions. Dulbecco’s Modified Eagle’s/Ham’s Nutrient Mixture F-12 (DMEF12) growth media (Genesee Scientific, San Diego, CA), horse serum (GE Healthcare Life Sciences, Logan, UT), fetal bovine serum (Thermo Fisher Scientific), insulin (Thermo Fisher Scientific, Waltham, MA), human recombinant EGF (Corning, Bedford, MA), hydrocortisone (BD Biosciences, Bedford, MA), cholera toxin (Enzo Life Sciences, Farmingdale, NY), penicillin/streptomycin (Thermo Fisher Sci.) and gentamycin (GE Healthcare Life Sciences) were used as previously described [12].

### Cell Treatments

N1-guanyl-1,7-diamineheptane (GC7), a spermidine analog that inhibits the activity of the hypusination enzyme deoxyhypusine synthase, was obtained from Biosearch Technologies (Petaluma, CA) and diluted in sterile deionized water. Fibronectin (Corning, Bedford, MA) coating/treatment was performed by pretreating plates or adding it to culture media at 10 ug/mL in sterile PBS at 37 °C for 1 hour (coating only). Control treatments used either sterile deionized water or PBS alone. UM7F8, a monoclonal anti-SLC3A2 (CD98hc) antibody with neutralizing effects, was gifted from our collaborator Dr. Joseph Cantor. The UM7F8 reagent was diluted in sterile phosphate buffer saline (PBS) and incubated with cells at 250 ng/mL for at least 48 hours in combination with an anti-mouse IgG secondary antibody from Jackson Immunoresearch (catalog #115-006-071) at 1 ug/mL. Control treatments used the anti-mouse IgG secondary antibody alone at 1 ug/mL. Palbociclib (PD0332991) was purchased from Tocris (catalog #47-861-0) and dissolved in deionized water at a stock concentration of 10 mg/mL. Cells were incubated with 1uM Palbociclib or control (i.e., sterile deionized water) beginning 24 hours after cell plating and throughout the course of time-lapse cell growth experiments.

### IncuCyte Spheroid Assay

Cells were seeded at 2,000 cells/well in spheroid media with or without fibronectin as previously described [58] using round-bottom ultra-low-attachment (ULA) 96-well plates. The IncuCyte SX1 system was used to collect bright-field images at 10X magnification every two hours. Largest object area was quantified using the Sartorius spheroid assay module. Time-course data over 48 hours were graphed and the area under the curve (AUC) was quantified using GraphPad Prism. Cells receiving pre-treatments were retreated upon plating.

### IncuCyte Cell Proliferation Assay

Cells were seeded at 3,000 cells/well in normal growth media onto uncoated or fibronectin 96-well plates. The IncuCyte SX1 system was used to collect phase-contrast images at 10X magnification every two hours. Cell confluence was quantified using the Sartorius basic cell analyzer assay module. Time-course data over five days were graphed and the area under the curve (AUC) was quantified using GraphPad Prism. Cells receiving pre-treatments were retreated upon plating. Palbociclib or control treatments were added 24 hours after cell seeding.

### Western Blot

Cells were plated on 6-well plates at 2e05 cells/well and one day later treated as indicated. Total protein lysates were collected using a standard RIPA buffer containing Pierce™ Protease and Phosphatase inhibitors (Thermo Fisher Sci.). Lysis was performed under rotation at 4 °C for three hours. Lysates were cleared by centrifugation at 12,000 RPM at 4 °C for 10 minutes. Protein concentrations were determined using the Coomassie-based Bradford Assay Reagent (Thermo Fisher Sci.) and absorbances were measured at 595 nm. 20 mg of total protein was loaded for each sample into NuPAGE™ 4-12% Bis-Tris gels (Thermo Fisher Sci.) and separated electrophoretically in NuPage LDS Running Buffer (Thermo Fisher Sci.) with 1 mM dithiothreitol (DTT). Spectra™ Multicolor Protein Ladder (Thermo Fisher Sci.) was loaded as protein size standards. Gels were then transferred using Pierce® Transfer Buffer (Thermo Fisher Sci.) onto 0.22 um Amersham™ Protran™ nitrocellulose membranes (GE Healthcare Life Science) at 30 V and 4 °C for 1 hour. Following transfer, total protein was visualized using PonceauS staining (Sigma-Aldrich, St. Louis, MO), blocked using 5% non-fat milk powder in tris-buffered saline with tween (TBST) and exposed to the following primary antibody solutions overnight at 4 °C: Hypusine (Thermo Fisher Sci.; 1:1000), eIF5A1/2 (Thermo Fisher Sci.; 1:1000), SLC3A2 (Abcam; 1:1000) and GAPDH (Prosci; 1:1000). HRP-conjugated goat anti-mouse/rabbit Pierce® secondary antibodies (Thermo Fisher Sci.) were diluted 1:10 000 in blocking solution and applied at room temperature for 1 hour. Protein bands were visualized using Pierce™ ECL Western Blotting Substrate kit (Thermo Fisher Sci.) and autoradiography film (Genesee Sci.) with exposures ranging from 10 sec to 30 min. Films were developed using a Medical Film Processor SRX-101 A (Konica Minolta, Inc., Tokyo, Japan) and scanned on an Officejet Pro 8600 (HP, Palo Alto, CA) for protein band quantification in Fiji (v.1.51n). Band densitometry quantification was performed using FIJI.

### Immunofluorescence & Structured Illumination Microscopy

Cells were plated as described above for Western blotting and treated with indicated cell treatment for 48 hrs. 48 hrs after incubation in treatment, cells were passaged and plated onto glass coverslips in 6-well plates containing either fibronectin (Corning. catalog #354008, 10 ug/mL/well) or no fibronectin (glass, 1 mL/well PBS) at a concentration of 1e5 cells/well. Cells were fixed in fresh 4% paraformaldehyde (Electron Microscopy Sciences, catalog #15713), permeabilized in 0.5% Triton-X100 (Fisher Scientific, catalog #BP151-100), treated with a 10% fetal bovine serum (FBS)/PBS blocking solution (Fisher bioreagents, catalog #BP9703-100) and stained for eIF5A1/2 polyclonal antibody (Thermo Fisher Sci. catalog #PA529204, 1:200) or SMAD3 (Cell Signaling Technology catalog #9523S, 1:100) in 2% BSA/PBS solution and the appropriate fluorescently-conjugated secondary antibody (R&D Systems catalog #NL006, 1:200). Coverslips were mounted onto slide with DAPI-containing ProLong Gold media. Structured illumination images were captured at 20X magnification using the Zeiss AXIO Imager.M2 microscope with Apotome.2 and Axiocam 506 monochorme camera.

Quantification of cytoplasmic and nuclear eIF5A1/2 and SMAD3 localization was performed with the cell image analysis software program Cell Profiler (Version 4.1.3). Standardization of protein staining intensities used a macro pipeline to ensure consistent settings/quantifications among different cell treatments. Settings included image illumination corrections, identification of primary objects (nuclei staining intensity settings), secondary objects (cell staining intensity settings), tertiary objects (cytoplasmic staining intensity settings), object intensity measurements, colocalization measurements, and the calculation of the ratio of cytoplasmic to nuclear intensity staining. Upon finalization of macro pipeline settings, images were analyzed and cytoplasmic to nuclear staining values were graphed and analyzed using GraphPad Prism.

### Flow Cytometry Analysis of Cell Cycle

Cells were plated as described above for Western blotting assays into uncoated or fibronectin-coated 6-well plates and treated with indicated cell treatments for 48 hrs. 48 hrs after incubation in treatment, cells were washed in PBS and trypsinized in 0.25% trypsin (Thermo Fisher Scientific, catalog# 15090046 1:10 with PBS) followed by centrifugation at 1000rpm for 5 min. Cells were fixed in ice-cold 70% ethanol/PBS on ice for 15 min.Cells pellets were incubated in 500μL propidium iodide (PI) solution containing PI, RNAse and 0.5% Triton-X100 for 40 minutes at 37 °C. Three mL PBS was added to tube and centrifuged at 1500 rpm for 5 minutes. Cell pellets were suspended in 500μL PBS and analyzed for side/forward scatter and PI fluorescence intensity a BD FACS Calibur. FlowJo analysis software was used to gate single-live cells and map the cell cycle profile to PI histograms.

### Bioinformatics

Literature-curated lists of genes implicated in the eIF5A1/2 hypusination pathway and intratumoral heterogeneity were assembled using PubMed. The resulting gene lists were then evaluated for biochemical and/or functional connections using the Cytoscape Agilent Literature Search. The resulting interactome was assessed for gene ontology enrichments using the DAVID database. Clinical implications associated with genomic or transcriptomic changes in the interactome nodes that made direct connections to either eIF5A1 or eIF5A2 genes were analyzed in over 2000 breast cancer patients using the METABRIC study in cBioPortal [59; 60]. Using this same cBioPortal data set, the relationship between genomic alteration of intratumoral heterogeneity transcription factors that interact with eIF5A1 or eIF5A2 and the mRNA expression levels of eIF5A1/2 hypusination pathway markers that independently predicted unfavorable patient survival was evaluated. Finally, the KM-Plotter database was used to evaluate overall survival for breast cancer patients across the four primary PAM50 subtypes in relationship to mRNA expression levels of DHPS and/or SLC3A2. Data were plotted and analyzed in Kaplan-Meier format using GraphPad Prism.

### Protein Analysis in Patient Samples

Protein products of the genes associated with eIF5A1/2 in the Cytoscape interactome were analyzed in human tissues using the Human Protein Atlas Tissue Atlas and Pathology Atlas resources for normal and cancerous breast cancer patient tissue, respectively [61; 62; 63]. The protein intensity staining levels for each antibody against each antigen were averaged and plotted using GraphPad Prism. Staining intensity assignments were made based on the Human Protein Atlas scoring scheme of 0-3, where 0 = not detected or N/A, 1 = low, 2 = medium, and 3 = high expression levels. The antibody average across all tissues were represented with a single dot in the graphs for each gene. Images of immunohistochemistry stained normal and tumor tissue sections were obtained from the Human Protein Atlas using indicated patients.

## ACKNOWLEDGEMENTS

We thank members of the Developmental Oncogene Laboratory (Kelber Lab) for helpful feedback throughout the duration of this project and constructive comments during manuscript preparation. This work was supported by California State University Northridge College of Science and Mathematics; Sidney Stern Memorial Trust; Sutter family; Aylozyan Family Foundation; NIH NIGMS grant SC1GM121182 (to J.A.K.); NIGMS grant SC1GM121182-07S1 (to J.A.K for G.O.S.); and NIGMS grant T34GM136450 (to M.E.Z. for J.J.M. and R.T.).

## REFERENCES

[1] A.C. Garrido-Castro, N.U. Lin, and K. Polyak, Insights into Molecular Classifications of Triple-Negative Breast Cancer: Improving Patient Selection for Treatment. Cancer Discov 9 (2019) 176–198.

[2] A.G. Waks, and E.P. Winer, Breast Cancer Treatment: A Review. JAMA 321 (2019) 288–300.

[3] F.A. Urra, S. Fuentes-Retamal, C. Palominos, Y.A. Rodriguez-Lucart, C. Lopez-Torres, and R. Araya-Maturana, Extracellular Matrix Signals as Drivers of Mitochondrial Bioenergetics and Metabolic Plasticity of Cancer Cells During Metastasis. Front Cell Dev Biol 9 (2021) 751301.

[4] E.A. Pietila, J. Gonzalez-Molina, L. Moyano-Galceran, S. Jamalzadeh, K. Zhang, L. Lehtinen, S.P. Turunen, T.A. Martins, O. Gultekin, T. Lamminen, K. Kaipio, U. Joneborg, J. Hynninen, S. Hietanen, S. Grenman, R. Lehtonen, S. Hautaniemi, O. Carpen, J.W. Carlson, and K. Lehti, Co-evolution of matrisome and adaptive adhesion dynamics drives ovarian cancer chemoresistance. Nat Commun 12 (2021) 3904.

[5] L. Fattet, H.Y. Jung, M.W. Matsumoto, B.E. Aubol, A. Kumar, J.A. Adams, A.C. Chen, R.L. Sah, A.J. Engler, E.B. Pasquale, and J. Yang, Matrix Rigidity Controls Epithelial-Mesenchymal Plasticity and Tumor Metastasis via a Mechanoresponsive EPHA2/LYN Complex. Dev Cell 54 (2020) 302–316 e7.

[6] M.C. Jones, J.A. Askari, J.D. Humphries, and M.J. Humphries, Cell adhesion is regulated by CDK1 during the cell cycle. J Cell Biol 217 (2018) 3203–3218.

[7] J.G. Lock, M.C. Jones, J.A. Askari, X. Gong, A. Oddone, H. Olofsson, S. Goransson, M. Lakadamyali, M.J. Humphries, and S. Stromblad, Reticular adhesions are a distinct class of cell-matrix adhesions that mediate attachment during mitosis. Nat Cell Biol 20 (2018) 1290–1302.

[8] R.E. Gough, M.C. Jones, T. Zacharchenko, S. Le, M. Yu, G. Jacquemet, S.P. Muench, J. Yan, J.D. Humphries, C. Jorgensen, M.J. Humphries, and B.T. Goult, Talin mechanosensitivity is modulated by a direct interaction with cyclin-dependent kinase-1. J Biol Chem 297 (2021) 100837.

[9] A.H. Cheung, C.H. Hui, K.Y. Wong, X. Liu, B. Chen, W. Kang, and K.F. To, Out of the cycle: Impact of cell cycle aberrations on cancer metabolism and metastasis. Int J Cancer (2022).

[10] S.J. Baker, P.I. Poulikakos, H.Y. Irie, S. Parekh, and E.P. Reddy, CDK4: a master regulator of the cell cycle and its role in cancer. Genes Cancer 13 (2022) 21–45.

[11] K. Fujimura, T. Wright, J. Strnadel, S. Kaushal, C. Metildi, A.M. Lowy, M. Bouvet, J.A. Kelber, and R.L. Klemke, A hypusine-eIF5A-PEAK1 switch regulates the pathogenesis of pancreatic cancer. Cancer Res 74 (2014) 6671–81.

[12] R. Guth, Y. Adamian, C. Geller, J. Molnar, J. Maddela, L. Kutscher, K. Bhakta, K. Meade, S.L. Kim, M. Agajanian, and J.A. Kelber, DHPS-dependent hypusination of eIF5A1/2 is necessary for TGFbeta/fibronectin-induced breast cancer metastasis and associates with prognostically unfavorable genomic alterations in TP53. Biochem Biophys Res Commun 519 (2019) 838–845.

[13] A. Arruabarrena-Aristorena, A. Zabala-Letona, and A. Carracedo, Oil for the cancer engine: The cross-talk between oncogenic signaling and polyamine metabolism. Sci Adv 4 (2018) eaar2606.

[14] M.B. Mathews, and J.W. Hershey, The translation factor eIF5A and human cancer. Biochemica and Biophysica Acta 1849 (2015) 836–44.

[15] M.A. Muniz Lino, Y. Palacios-Rodriguez, S. Rodriguez-Cuevas, V. Bautista-Pina, L.A. Marchat, E. Ruiz-Garcia, H. Astudillo-de la Vega, A.E. Gonzalez-Santiago, A. Flores-Perez, J. Diaz-Chavez, A. Carlos-Reyes, E. Alvarez-Sanchez, and C. Lopez-Camarillo, Comparative proteomic profiling of triple-negative breast cancer reveals that up-regulation of RhoGDI-2 is associated to the inhibition of caspase 3 and caspase 9. J Proteomics 111 (2014) 198–211.

[16] S. Coni, S.M. Serrao, Z.N. Yurtsever, L. Di Magno, R. Bordone, C. Bertani, V. Licursi, Z. Ianniello, P. Infante, M. Moretti, M. Petroni, F. Guerrieri, A. Fatica, A. Macone, E. De Smaele, L. Di Marcotullio, G. Giannini, M. Maroder, E. Agostinelli, and G. Canettieri, Blockade of EIF5A hypusination limits colorectal cancer growth by inhibiting MYC elongation. Cell Death Dis 11 (2020) 1045.

[17] Y. Liu, R. Liu, P. Fu, F. Du, Y. Hong, M. Yao, X. Zhang, and S. Zheng, N1-Guanyl-1,7-Diaminoheptane Sensitizes Estrogen Receptor Negative Breast Cancer Cells to Doxorubicin by Preventing Epithelial-Mesenchymal Transition through Inhibition of Eukaryotic Translation Initiation Factor 5A2 Activation. Cellular Physiology and Biochemistry 36 (2015) 2494–503.

[18] M. Aksu, S. Trakhanov, and D. Gorlich, Structure of the exportin Xpo4 in complex with RanGTP and the hypusine-containing translation factor eIF5A. Nat Commun 7 (2016) 11952.

[19] C.C. Feral, A. Zijlstra, E. Tkachenko, G. Prager, M.L. Gardel, M. Slepak, and M.H. Ginsberg, CD98hc (SLC3A2) participates in fibronectin matrix assembly by mediating integrin signaling. J Cell Biol 178 (2007) 701–11.

[20] C. Wang, Z. Chen, L. Nie, M. Tang, X. Feng, D. Su, H. Zhang, Y. Xiong, J.M. Park, and J. Chen, Extracellular signal-regulated kinases associate with and phosphorylate DHPS to promote cell proliferation. Oncogenesis 9 (2020) 85.

[21] S. Cano-Crespo, J. Chillaron, A. Junza, G. Fernandez-Miranda, J. Garcia, C. Polte, R.d.l.B. L, Z. Ignatova, O. Yanes, A. Zorzano, C. Stephan-Otto Attolini, and M. Palacin, CD98hc (SLC3A2) sustains amino acid and nucleotide availability for cell cycle progression. Sci Rep 9 (2019) 14065.

[22] S. Yuan, R.J. Norgard, and B.Z. Stanger, Cellular Plasticity in Cancer. Cancer Discov (2019).

[23] A.E. Davies, and J.G. Albeck, Microenvironmental Signals and Biochemical Information Processing: Cooperative Determinants of Intratumoral Plasticity and Heterogeneity. Front Cell Dev Biol 6 (2018) 44.

[24] H. Hammerlindl, and H. Schaider, Tumor cell-intrinsic phenotypic plasticity facilitates adaptive cellular reprogramming driving acquired drug resistance. J Cell Commun Signal 12 (2018) 133–141.

[25] H. Sun, J. Zeng, Z. Miao, K.C. Lei, C. Huang, L. Hu, S.M. Su, U.I. Chan, K. Miao, X. Zhang, Zhang, S. Guo, S. Chen, Y. Meng, M. Deng, W. Hao, H. Lei, Y. Lin, Z. Yang, D. Tang, K.H. Wong, X.D. Zhang, X. Xu, and C.X. Deng, Dissecting the heterogeneity and tumorigenesis of BRCA1 deficient mammary tumors via single cell RNA sequencing. Theranostics 11 (2021) 9967–9987.

[26] D.L. Ellsworth, H.L. Blackburn, C.D. Shriver, S. Rabizadeh, P. Soon-Shiong, and R.E. Ellsworth, Single-cell sequencing and tumorigenesis: improved understanding of tumor evolution and metastasis. Clin Transl Med 6 (2017) 15.

[27] I. Novita Sari, T. Setiawan, K. Seock Kim, Y. Toni Wijaya, K. Won Cho, and H. Young Kwon, Metabolism and function of polyamines in cancer progression. Cancer Lett 519 (2021) 91–104.

[28] M.H. Park, and E.C. Wolff, Hypusine, a polyamine-derived amino acid critical for eukaryotic translation. J Biol Chem 293 (2018) 18710–18718.

[29] A. Mandal, S. Mandal, and M.H. Park, Global quantitative proteomics reveal up-regulation of endoplasmic reticulum stress response proteins upon depletion of eIF5A in HeLa cells. Sci Rep 6 (2016) 25795.

[30] J. Liang, and Z. Sun, Overexpression of membranal SLC3A2 regulates the proliferation of oral squamous cancer cells and affects the prognosis of oral cancer patients. J Oral Pathol Med 50 (2021) 371–377.

[31] L.D. Gamble, S. Purgato, J. Murray, L. Xiao, D.M.T. Yu, K.M. Hanssen, F.M. Giorgi, D.R. Carter, A.J. Gifford, E. Valli, G. Milazzo, A. Kamili, C. Mayoh, B. Liu, G. Eden, S. Sarraf, S. Allan, S. Di Giacomo, C.L. Flemming, A.J. Russell, B.B. Cheung, A. Oberthuer, W.B. London, M. Fischer, T.N. Trahair, J.I. Fletcher, G.M. Marshall, D.S. Ziegler, M.D. Hogarty, M.R. Burns, G. Perini, M.D. Norris, and M. Haber, Inhibition of polyamine synthesis and uptake reduces tumor progression and prolongs survival in mouse models of neuroblastoma. Sci Transl Med 11 (2019).

[32] T.C.J. Tan, V. Kelly, X. Zou, D. Wright, T. Ly, and R. Zamoyska, Translation factor eIF5a is essential for IFNgamma production and cell cycle regulation in primary CD8(+) T lymphocytes. Nat Commun 13 (2022) 7796.

[33] G.W. Prager, C.C. Feral, C. Kim, J. Han, and M.H. Ginsberg, CD98hc (SLC3A2) interaction with the integrin beta subunit cytoplasmic domain mediates adhesive signaling. J Biol Chem 282 (2007) 24477–84.

[34] C.C. Feral, N. Nishiya, C.A. Fenczik, H. Stuhlmann, M. Slepak, and M.H. Ginsberg, CD98hc (SLC3A2) mediates integrin signaling. Proc Natl Acad Sci U S A 102 (2005) 355–60.

[35] R. El Ansari, M.L. Craze, M. Diez-Rodriguez, C.C. Nolan, I.O. Ellis, E.A. Rakha, and A.R. Green, The multifunctional solute carrier 3A2 (SLC3A2) confers a poor prognosis in the highly proliferative breast cancer subtypes. Br J Cancer 118 (2018) 1115–1122.

[36] J.C. Montero, E. Calvo-Jimenez, S. Del Carmen, M. Abad, A. Ocana, and A. Pandiella, Surfaceome analyses uncover CD98hc as an antibody drug-conjugate target in triple negative breast cancer. J Exp Clin Cancer Res 41 (2022) 106.

[37] J.N. Ablack, J. Ortiz, J. Bajaj, K. Trinh, F. Lagarrigue, J.M. Cantor, T. Reya, and M.H. Ginsberg, MARCH Proteins Mediate Responses to Antitumor Antibodies. J Immunol 205 (2020) 2883–2892.

[38] J.M. Cantor, and M.H. Ginsberg, CD98 at the crossroads of adaptive immunity and cancer. J Cell Sci 125 (2012) 1373–82.

[39] R. Yan, X. Zhao, J. Lei, and Q. Zhou, Structure of the human LAT1-4F2hc heteromeric amino acid transporter complex. Nature 568 (2019) 127–130.

[40] E. Boulter, S. Estrach, F.S. Tissot, M.L. Hennrich, L. Tosello, L. Cailleteau, L.R. de la Ballina, S. Pisano, A.C. Gavin, and C.C. Feral, Cell metabolism regulates integrin mechanosensing via an SLC3A2-dependent sphingolipid biosynthesis pathway. Nat Commun 9 (2018) 4862.

[41] M. Tauc, M. Cougnon, R. Carcy, N. Melis, T. Hauet, L. Pellerin, N. Blondeau, and D.F. Pisani, The eukaryotic initiation factor 5A (eIF5A1), the molecule, mechanisms and recent insights into the pathophysiological roles. Cell Biosci 11 (2021) 219.

[42] B. Maier, T. Ogihara, A.P. Trace, S.A. Tersey, R.D. Robbins, S.K. Chakrabarti, C.S. Nunemaker, N.D. Stull, C.A. Taylor, J.E. Thompson, R.S. Dondero, E.C. Lewis, C.A. Dinarello, J.L. Nadler, and R.G. Mirmira, The unique hypusine modification of eIF5A promotes islet beta cell inflammation and dysfunction in mice. J Clin Invest 120 (2010) 2156–70.

[43] S.B. Lee, J.H. Park, J. Kaevel, M. Sramkova, R. Weigert, and M.H. Park, The effect of hypusine modification on the intracellular localization of eIF5A. Biochem Biophys Res Commun 383 (2009) 497–502.

[44] M. Ishfaq, K. Maeta, S. Maeda, T. Natsume, A. Ito, and M. Yoshida, The role of acetylation in the subcellular localization of an oncogenic isoform of translation factor eIF5A. Biosci Biotechnol Biochem 76 (2012) 2165–7.

[45] M.K. Hayward, J.M. Muncie, and V.M. Weaver, Tissue mechanics in stem cell fate, development, and cancer. Dev Cell 56 (2021) 1833–1847.

[46] P. Moreno-Layseca, and C.H. Streuli, Signalling pathways linking integrins with cell cycle progression. Matrix Biol 34 (2014) 144–53.

[47] E. Gutierrez, B.S. Shin, C.J. Woolstenhulme, J.R. Kim, P. Saini, A.R. Buskirk, and T.E. Dever, eIF5A promotes translation of polyproline motifs. Mol Cell 51 (2013) 35–45.

[48] A.P. Schuller, C.C. Wu, T.E. Dever, A.R. Buskirk, and R. Green, eIF5A Functions Globally in Translation Elongation and Termination. Mol Cell 66 (2017) 194–205 e5.

[49] V. Pelechano, and P. Alepuz, eIF5A facilitates translation termination globally and promotes the elongation of many non polyproline-specific tripeptide sequences. Nucleic Acids Res 45 (2017) 7326–7338.

[50] M. Agajanian, A. Campeau, M. Hoover, A. Hou, D. Brambilla, S.L. Kim, R.L. Klemke, and J.A. Kelber, PEAK1 Acts as a Molecular Switch to Regulate Context-Dependent TGFbeta Responses in Breast Cancer. PLoS One 10 (2015) e0135748.

[51] C. Gorrini, F. Loreni, V. Gandin, L.A. Sala, N. Sonenberg, P.C. Marchisio, and S. Biffo, Fibronectin controls cap-dependent translation through beta1 integrin and eukaryotic initiation factors 4 and 2 coordinated pathways. Proc Natl Acad Sci U S A 102 (2005) 9200–5.

[52] J.D. Humphries, A. Byron, and M.J. Humphries, Integrin ligands at a glance. J Cell Sci 119 (2006) 3901–3.

[53] V. Rapisarda, M. Borghesan, V. Miguela, V. Encheva, A.P. Snijders, A. Lujambio, and A. O’Loghlen, Integrin Beta 3 Regulates Cellular Senescence by Activating the TGF-beta Pathway. Cell Rep 18 (2017) 2480–2493.

[54] M. Alvarez-Fernandez, and M. Malumbres, Mechanisms of Sensitivity and Resistance to CDK4/6 Inhibition. Cancer Cell 37 (2020) 514–529.

[55] A. Yoshida, Y. Bu, S. Qie, J. Wrangle, E.R. Camp, E.S. Hazard, G. Hardiman, R. de Leeuw, K.E. Knudsen, and J.A. Diehl, SLC36A1-mTORC1 signaling drives acquired resistance to CDK4/6 inhibitors. Sci Adv 5 (2019) eaax6352.

[56] J. Kahlhofer, and D. Teis, The human LAT1-4F2hc (SLC7A5-SLC3A2) transporter complex: Physiological and pathophysiological implications. Basic Clin Pharmacol Toxicol (2022).

[57] M.C. Jones, J. Zha, and M.J. Humphries, Connections between the cell cycle, cell adhesion and the cytoskeleton. Philos Trans R Soc Lond B Biol Sci 374 (2019) 20180227.

[58] J.A. Kelber, T. Reno, S. Kaushal, C. Metildi, T. Wright, K. Stoletov, J.M. Weems, F.D. Park, E. Mose, Y. Wang, R.M. Hoffman, A.M. Lowy, M. Bouvet, and R.L. Klemke, KRas induces a Src/PEAK1/ErbB2 kinase amplification loop that drives metastatic growth and therapy resistance in pancreatic cancer. Cancer Res 72 (2012) 2554–64.

[59] C. Curtis, S.P. Shah, S.F. Chin, G. Turashvili, O.M. Rueda, M.J. Dunning, D. Speed, A.G. Lynch, S. Samarajiwa, Y. Yuan, S. Graf, G. Ha, G. Haffari, A. Bashashati, R. Russell, S. McKinney, M. Group, A. Langerod, A. Green, E. Provenzano, G. Wishart, S. Pinder, P. Watson, F. Markowetz, L. Murphy, I. Ellis, A. Purushotham, A.L. Borresen-Dale, J.D. Brenton, S. Tavare, C. Caldas, and S. Aparicio, The genomic and transcriptomic architecture of 2,000 breast tumours reveals novel subgroups. Nature 486 (2012) 346–52.

[60] O.M. Rueda, S.J. Sammut, J.A. Seoane, S.F. Chin, J.L. Caswell-Jin, M. Callari, R. Batra, B. Pereira, A. Bruna, H.R. Ali, E. Provenzano, B. Liu, M. Parisien, C. Gillett, S. McKinney, A.R. Green, L. Murphy, A. Purushotham, I.O. Ellis, P.D. Pharoah, C. Rueda, S. Aparicio,C. Caldas, and C. Curtis, Dynamics of breast-cancer relapse reveal late-recurring ER-positive genomic subgroups. Nature 567 (2019) 399–404.

[61] M. Uhlen, C. Zhang, S. Lee, E. Sjostedt, L. Fagerberg, G. Bidkhori, R. Benfeitas, M. Arif, Z. Liu, F. Edfors, K. Sanli, K. von Feilitzen, P. Oksvold, E. Lundberg, S. Hober, P. Nilsson, J. Mattsson, J.M. Schwenk, H. Brunnstrom, B. Glimelius, T. Sjoblom, P.H. Edqvist, D. Djureinovic, P. Micke, C. Lindskog, A. Mardinoglu, and F. Ponten, A pathology atlas of the human cancer transcriptome. Science 357 (2017).

[62] M. Uhlen, L. Fagerberg, B.M. Hallstrom, C. Lindskog, P. Oksvold, A. Mardinoglu, A. Sivertsson, C. Kampf, E. Sjostedt, A. Asplund, I. Olsson, K. Edlund, E. Lundberg, S. Navani, C.A. Szigyarto, J. Odeberg, D. Djureinovic, J.O. Takanen, S. Hober, T. Alm, P.H. Edqvist, H. Berling, H. Tegel, J. Mulder, J. Rockberg, P. Nilsson, J.M. Schwenk, M. Hamsten, K. von Feilitzen, M. Forsberg, L. Persson, F. Johansson, M. Zwahlen, G. von Heijne, J. Nielsen, and F. Ponten, Proteomics. Tissue-based map of the human proteome. Science 347 (2015) 1260419.

[63] M. Uhlen, P. Oksvold, L. Fagerberg, E. Lundberg, K. Jonasson, M. Forsberg, M. Zwahlen, C. Kampf, K. Wester, S. Hober, H. Wernerus, L. Bjorling, and F. Ponten, Towards a knowledge-based Human Protein Atlas. Nat Biotechnol 28 (2010) 1248–50.

