## Supplemental Figure 1 for "Fibronectin, DHPS and SLC3A2 Signaling Cooperate to Control Tumor Spheroid Growth, Subcellular eIF5A1/2 Distribution and CDK4/6 Inhibitor Resistance"

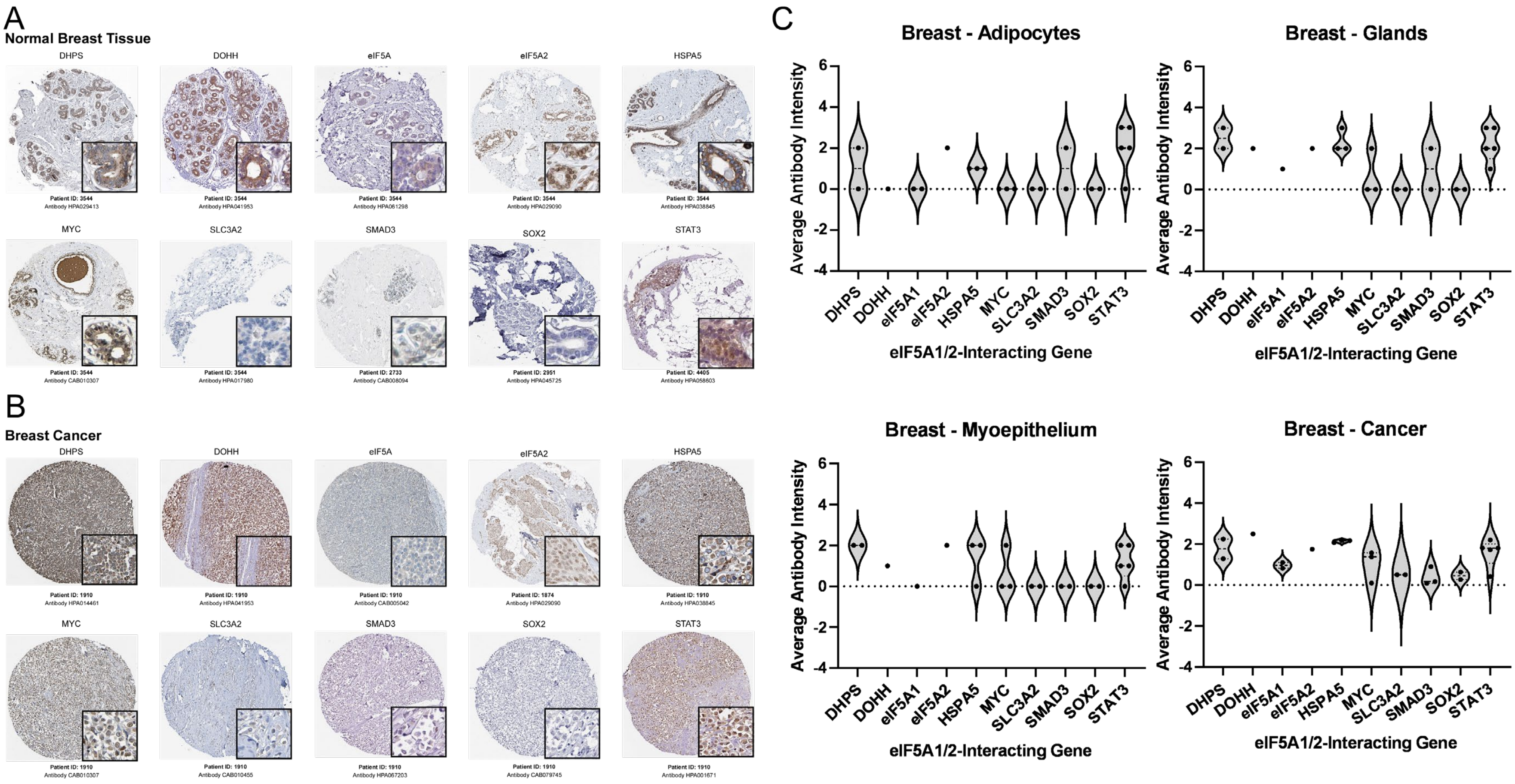

**Supplemental Figure 1: eIF5A1/2-Interacting genes display differential staining patterns across benign and malignant breast tissue.** A-B. Representative tissue images from normal (A) and malignant (B) breast tissue. C. Quantification of average antibody intensity as a function of antibody for each of the eIF5A1/2-Interacting Genes across non-cancerous breast cell types and breast cancer samples. Numerical value of 0-3 were assigned based on antibody intensity staining (0 = not detected or N/A, 1 = low, 2 = medium, 3 = high) as reported in the Human Protein Atlas database. Each dot in graphs represents different antibody for respective gene.
