## Supplemental Figure 2 for "Fibronectin, DHPS and SLC3A2 Signaling Cooperate to Control Tumor Spheroid Growth, Subcellular eIF5A1/2 Distribution and CDK4/6 Inhibitor Resistance"

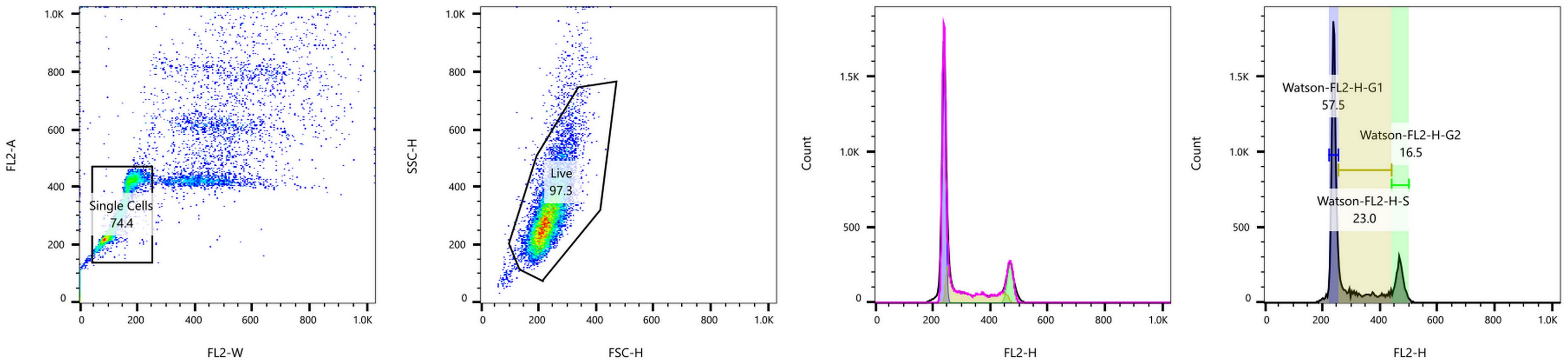

**Supplemental Figure 2: Gating scheme for analysis of cell cycle data using flow cytometry. Data were analyzed using FlowJo and the Watson (Pragmatic) model within the cell cycle analysis application.**
